# Tracking Real-Time Changes in Working Memory Updating and Gating with the Event-Based Eye-Blink Rate

**DOI:** 10.1101/098103

**Authors:** Rachel Rac-Lubashevsky, Heleen A. Slagter, Yoav Kessler

**Author notes:** Corresponding author: Rachel Rac-Lubashevsky, Department of Brain and Cognitive Sciences and Zlotowski Center for Neuroscience, Ben-Gurion University of the Negev, POB 653, Beer-Sheva, Israel, 84105.

## Abstract

Effective working memory (WM) functioning depends on the gating process that regulates the balance between maintenance and updating of WM. The present study used the event-based eye-blink rate (ebEBR), which presumably reflects phasic striatal dopamine activity, to examine how the cognitive processes of gating and updating separately facilitate flexible updating of WM contents and the potential involvement of dopamine in these processes. Realtime changes in eye blinks were tracked during performance on the reference-back task, in which demands on these two processes were independently manipulated. In all three experiments, trials that required WM updating and trials that required gate switching were both associated with increased ebEBR. These results may support the *prefrontal cortex basal ganglia WM model* (PBWM) by linking updating and gating to striatal dopaminergic activity. In Experiment 3, the ebEBR was used to determine what triggers gate switching. We found that switching to an updating mode (gate opening) was more stimulus driven and retroactive than switching to a maintenance mode, which was more context driven. Together, these findings show that the ebEBR – an inexpensive, non-invasive, easy-to-use measure – can be used to track changes in WM demands during task performance and, hence, possibly striatal dopamine activity.

## Introduction

Working memory (WM) is a dynamic system in which goal-relevant information is temporarily maintained and manipulated. Effective WM functioning depends on filtering or ‘gating’ mechanisms that “decide” which information is relevant and, hence, should enter into WM, and which is not and should be left outside. Optimizing the balance between these two requirements is described in the literature as the stability-flexibility dilemma ^1–3^. WM faces two seemingly opposing demands. First, the maintained information within WM must be shielded from being overridden by task-irrelevant input. Due to its capacity limit ^4^, storage of irrelevant information in WM would necessarily be at the expense of the ability to maintain relevant content. At the same time, task-relevant information must be recognized and entered into WM to accommodate relevant changes in the environment. Therefore, encoding information to WM must be selective^5–8^.

The prefrontal cortex basal ganglia WM model (PBWM) ^9,10^ (for alternative models see ^11,12^) is a physiologically based computational model that describes the mechanisms by which maintenance and updating are coordinated. According to the PBWM, the prefrontal cortex (PFC) is responsible for robust, active maintenance of information in a distractor-resistant manner, achieved through recurrent excitation ^10,11,13–17^ Controlling the flow of information into WM is achieved by the basal ganglia (BG), which serves as a “gate” to WM ^6,18^. When closed, the gate enables active maintenance and stability within WM by preventing irrelevant input from entering WM. However, when new input is relevant, the gate opens and enables flexible updating of the existing representations. Control over the state of the gate (namely, open or closed) is mediated through dopaminergic (DA) activity ^9,11,20,21^. The PBWM model assumes that the gate to WM is closed by default, and, hence, each instance of updating should be followed by gate closing.

In line with the PBWM model, a large body of work has shown that DA serves an important role both in supporting the maintenance of and updating WM as well as in their coordination. The stability of representation in WM is managed by tonic DA activity in the PFC, whereas phasic DA from the dorsal striatum drives the gate opening signal that leads to the updating of WM by disinhibiting the thalamus. However, the effectiveness of the phasic DA to override the tonic DA signal depends on the initial striatal tonic level ^12,22,23^. The disinhibition of the thalamus gates information into the PFC and thereby flexibly updates WM. This updating is also marked by the phasic DA signal ^10,24^.

A growing body of research indicates that spontaneous eye blinks (sEBs) may be an effective measure of striatal DA activity. sEBs are endogenous and unconscious responses^25^ that occur in the absence of any evident stimulus. Although the neural mechanisms that underlie sEBs are not yet fully understood, converging evidence from clinical and pharmacological studies indicates a positive correlation between striatal DA activity and the rate of eye blinks (i.e., the number of sEBs per minute) (sEBR) in the resting state ^26–30^. For example, two disorders characterized by DA dysfunction, Parkinson’s disease and schizophrenia, are associated with decreased ^28,29^ and increased ^27,32^ sEBR, respectively. Furthermore, DA agonists increase the sEBR, whereas DA antagonists decrease the sEBR^30,33–35^. Notably, the sEBR in resting-state conditions is correlated with subsequent performance of cognitive tasks that are known to depend on DA neurotransmission, including the stop-signal task^36^, attentional blink ^37^, attentional bias^38^, and task switching^39^. As these associations were found using resting-state sEBR, they suggest a relationship between tonic DA activity and cognitive functioning. Moreover, the sEBR has been related to avoidance learning and not to positive learning, which may suggest that the sEBR specifically reflects the activity of the DA D2 receptor^40^. This idea is supported by a recent PET study in monkeys, which found a strong correlation between the sEBR and D2-like receptor availability in the ventral striatum and caudate nucleus^41^. Furthermore, in this study, D2-like receptor availability correlated with D2-like receptor agonist-induced changes in the sEBR and the density of D2-like receptors determined in vitro. Thus, convergent evidence from different lines of research indicates that striatal DA activity regulates the sEBR.

sEBR provide an easy-to-obtain, non-invasive and inexpensive method for assessing the relationship between striatal DA function and behavior without the need to alter the natural DA activity in the brain. However, previous work has left unresolved to what extent the sEBR can be used to track real-time changes in DA activity *during* task performance as a function of task demands. Task-evoked eye blinks is a relatively new method^42–49^ compared to the resting-state method. To the best of our knowledge, only two studies, both in infants, have used event-based eye-blink rate (ebEBR) to test the involvement of fronto-striatal DA in WM updating. The first was conducted during an incidental hierarchical rule-learning task in 8-month-old infants^49^. The authors found increased ebEBR when the task rule was updated in WM compared to that observed when it was repeated. The second was conducted during an A-not-B WM task in 10-month-old infants^42^. The authors found that the ebEBR increased when the location of the hidden toy had to be updated in WM compared to that observed when the location of the toy was revealed. These initial findings indicate that the ebEBR is dynamically modulated by WM processes known to depend on DA activity, although it is unclear to what extent these findings extend to the adult brain.

The aim of the current study was to shed light on the involvement of DA in gating and WM updating by examining task demand-related changes in eye-blink rate during performance in the reference-back task^50,51^. The reference-back task is a novel paradigm that allows for separation of processes related to WM updating from processes related to gate opening and closing. This task is composed of two types of trials, *reference* and *comparison,* which are indicated by different colors (e.g., a red or blue frame surrounding the stimulus, respectively; see Fig. 1). In each trial, participants are required to indicate whether the presented stimulus (’X’ or ‘O’) is the same as or different from the most recent stimulus that appeared within a red frame, namely the “reference” stimulus. Accordingly, each trial in this task requires comparing the presented stimulus and the reference stimulus. While comparison trials (in which the stimulus was presented inside a blue frame) only require a same/different decision, reference trials (in which the stimulus was presented inside a red frame) in addition, require one to update WM with the presented stimulus. This is because each reference stimulus would serve as a reference to which the following trials would be compared. Thus, reference trials require opening the gate to WM to enable updating. By contrast, comparison trials do not require WM updating. Instead, these trials require one to continue maintaining the last reference stimulus in WM. Because each comparison trial is also compared to the last reference trial, the reference needs to be protected from being overwritten by changes in comparison trials. Hence, the gate over WM should be closed in these trials. Previous results using this paradigm^50,51^ have demonstrated that (a) performance in reference trials is slower than in comparison trials, supporting the additional updating process required in the former, and (b) switching between the two trial-types is associated with an additional cost, reflecting the time taken to open or close the gate to WM^52,53^.

**Figure 1.**
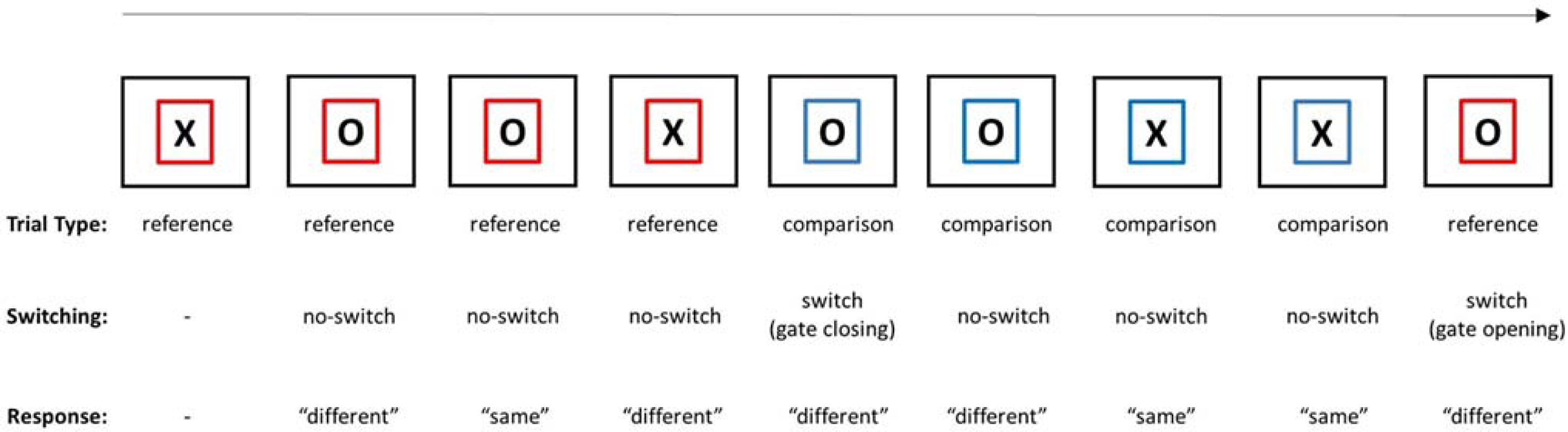
The reference-back task. Trials with a red frame are reference trials, and trials with a blue frame are comparison trials (see main text for details). The sequence length for each trial type was a constant of 4 trials. There was a fixation display after the response for 2-sec, thus creating a 2-sec inter-trial interval (ITI). The state of the gate and the correct response for each trial are indicated below each stimulus display.

In three experiments, we intended to determine whether the ebEBR can be used to track changes in demands on gating and updating. Inspired by the PBWM model and our previous results^50,51^ and under the assumption that the ebEBR reflects phasic DA activity, we predicted that WM updating and gate opening would be associated with an increase in the ebEBR. Gate closing would not be accompanied by an increase in the ebEBR because, as implied in the model, gate closing is the default state of the gate and does not require phasic DA. In Experiment 1, we tested this main prediction. In Experiment 2, we aimed to replicate the findings of Experiment 1 and extend these by examining the optimal window size in which ebEBR is sensitive to updating and gating. Finally, in Experiment 3, we aimed at testing the conditions required to open and close the gate. Specifically, we aimed to determine whether it is possible to prepare for gate opening and closing before the stimulus is presented. This was tested by cuing the condition, by presenting a colored frame (indicating the trial-type) for 4-sec prior to the presentation of the probe (X or O). Finding an ebEBR effect during the cuing interval would indicate that gating is context-driven. By contrast, if the ebEBR effect is only observed after the stimulus was presented, this result could indicate that gating is stimulus-driven. Stimulus-driven gating is a retroactive strategy, whereby all of the information on the input is required to make a decision to switch the state or not. Alternatively, context-driven gating would result from a more proactive strategy, whereby the stimulus is not a crucial element of the decision, but rather, the context is sufficient.

## Results

### Experiment 1

Our main prediction was that changes in WM task demands, specifically gate opening and updating, would be associated with changes in the ebEBR. A two-way ANOVA was conducted on the ebEBR data with Trial-Type (reference, comparison) and Switching (switch, no-switch) as the within-subject independent variables (see Fig. 2). Indeed, ebERB was significantly higher in the reference than in the comparison trials, *F(1,18)*=*13.63, MSe*=*.0013, p*=*.002, η_p_^2^*=*.43.* The ebEBR was also significantly higher in the switch compared to the no-switch trials (this difference will be referred as the switching effect from here on), *F(1,18)*=*24.48, MSe*=*.0015, p*<*.001, η_p_^2^*=*.58.* The two-way interaction was not significant, *F(1,18)*=*.37, MSe*=*.0010, p*=*.55, η_p_^2^*=*.02.*

**Figure 2.**
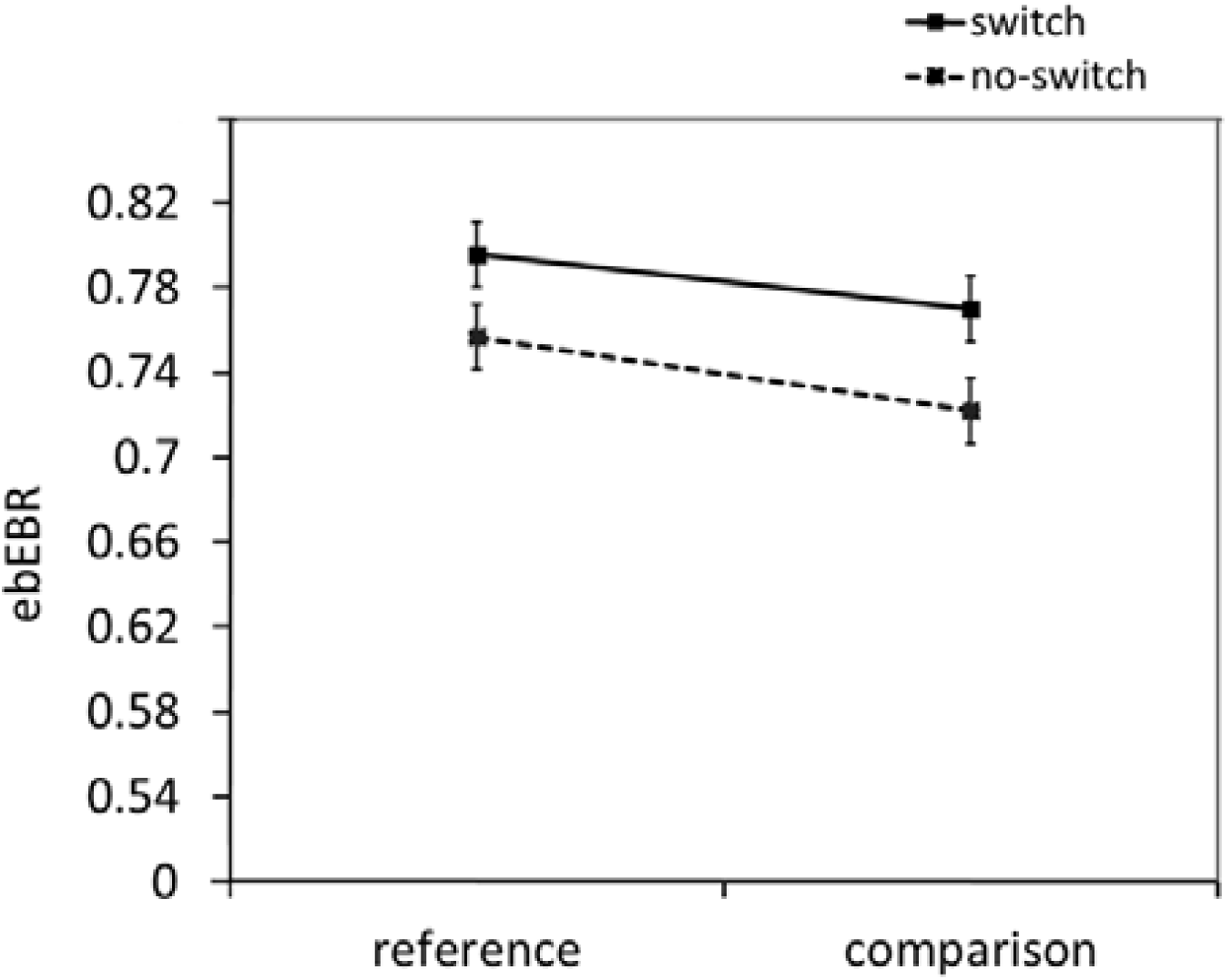
The ebEBR results of Experiment 1. Error bars represent 95% confidence intervals^59^.

### Experiment 2

To test the optimal window size in which the ebEBR is sensitive to task demands, we divided the 4-sec time window after stimulus presentation, into two halves of 2-sec each. A three-way ANOVA was conducted on the ebEBR data with Segment Part (first half, second half), Trial-Type (reference, comparison) and Switching (switch, no-switch) as within-subject independent variables (see Fig. 3). As before, the ebERB was significantly higher in the reference than in the comparison trials, *F(1,19)*=*15.37, MSe*=*.0015, p*<*.001, η_p_^2^*=*.45,* and the switching effect was also significant, *F(1,19)*=*11.41, MSe*=*.0015, p*=*.003, η_p_^2^*=*.37.* The twoway interaction between Switching and Segment Part was significant as well, *F(1,19)*=*4.91, MSe*=*.0015, p*=*.04, η_p_^2^*=*.47* (see Fig. 3). Specifically, the simple main effect of Switching was significant in the second time window, *F(1,19)*=*16.57, MSe*=*.0015, p*<*.001, η_p_^2^*=*.20,* but not in the first, *F(1,19)*=*.62, MSe*=*.0016, p*=*.44, η_p_^2^*=*.03.* Thus, the switching effect was more pronounced in the second than in the first half of the trial. At this stage, the reasons for the delayed effect of switching in Experiment 2 compared to that in Experiment 1 are not fully understood. One possibility is that the prolonged ITI in Experiment 2 led to the adoption of a more retroactive strategy, in which updating and possibly gating were carried out after the response was indicated. The two-way interaction between Segment Part and Trial-Type was marginally significant, *F(1,19)*=*4.20, MSe*=*.0013, p*=*.05, η_p_^2^*=*.18.* The three-way interaction was not significant, *F(1,19)*=*1.20, MSe*=*.0012, P*=*0.29, η_p_^2^*=*.06.*

**Figure 3.**
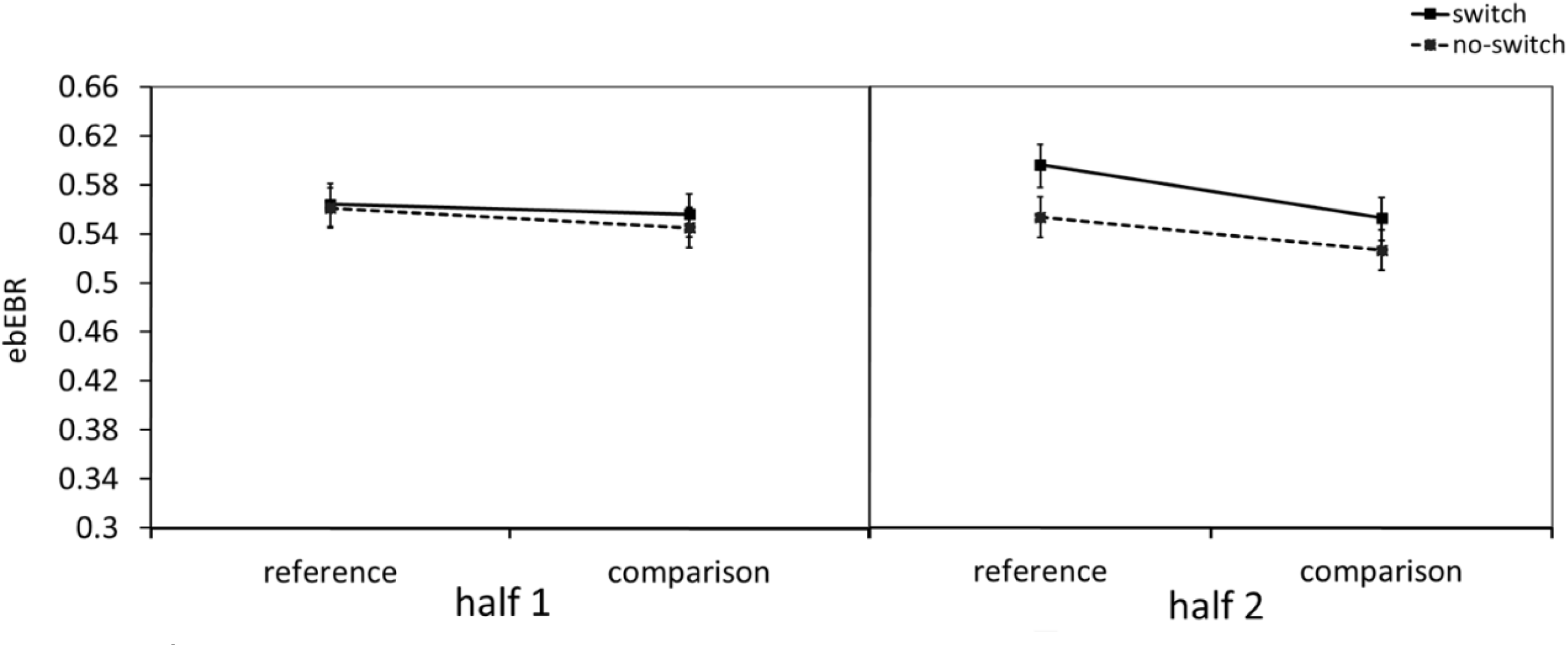
The ebEBR results of Experiment 2. The 4-sec segment length was divided into two parts, 2 sec each. The ebEBR as a function of Segment part, Trial Type and Switching is presented. The effect of switching was observed only in the second half of the segment. Error bars represent 95% confidence intervals.

### Experiment 3

To test what triggers the gate, a “random version” of the reference-back task was used in which both trial-types could appear in each trial with equal probabilities (see Fig. 5). Crucially, each trial began with a 4-sec of cue, which indicated the upcoming trial-type before the stimulus was presented but did not provide explicit information about the stimulus identity. Thus, in the cue phase, participants knew if this would be a reference or a comparison trial, which also implies whether this trial would be a switch or a no-switch trial. However, as they had no information about the stimulus identity at this point, they could not make response-related decisions and could not prepare a motor response. If a switching effect in the ebEBR was observed in the cue phase, this result would suggest that the processes involved in switch trials (presumably gate opening and closing) do not require the stimulus. However, if a switching effect was detected in the ebEBR only after the stimulus was presented, this result would support the stimulus-driven hypothesis.

## Cue phase analysis

A three-way ANOVA was conducted on the ebEBR data with Segment Part (first half, second half), Trial-Type (reference, comparison) and Switching (switch, no-switch) as the within-subject independent variables (see Fig. 4a). The effect of Segment Part was significant, *F(1,20)*=*79.74, MSe*=*.0129, p*<*.001, η_p_^2^*=*.80,* as was the two-way interaction between Segment Part and Switching, *F(1,20)*=*5.05, MSe*=*.0019, p*=*.04, η_p_^2^*=*.20.* This interaction effect indicated a larger effect for switching, reflected by the larger difference in the ebEBR between switch and no-switch trials, in the first half of the segment than in the second half (0.03 vs. −0.003 ebEBR per second, respectively). The two-way interaction between Switching and Trial-Type was also significant, *F(1,20)*=*11.89, MSe*=*.0020, p*=*.002, η_p_^2^*=*.37,* reflecting a significant switching effect only in comparison trials, *F(1,20)*=*13.72, MSe*=*.002, p*=*.001, η_p_^2^*=*.41,* but not in reference trials, *F(1,20)*=*1.90, MSe*=*.0016, p*=*.18, η_p_^2^*=*.09.* None of the other effects were significant, all Fs<3.61.

**Figure 4.**
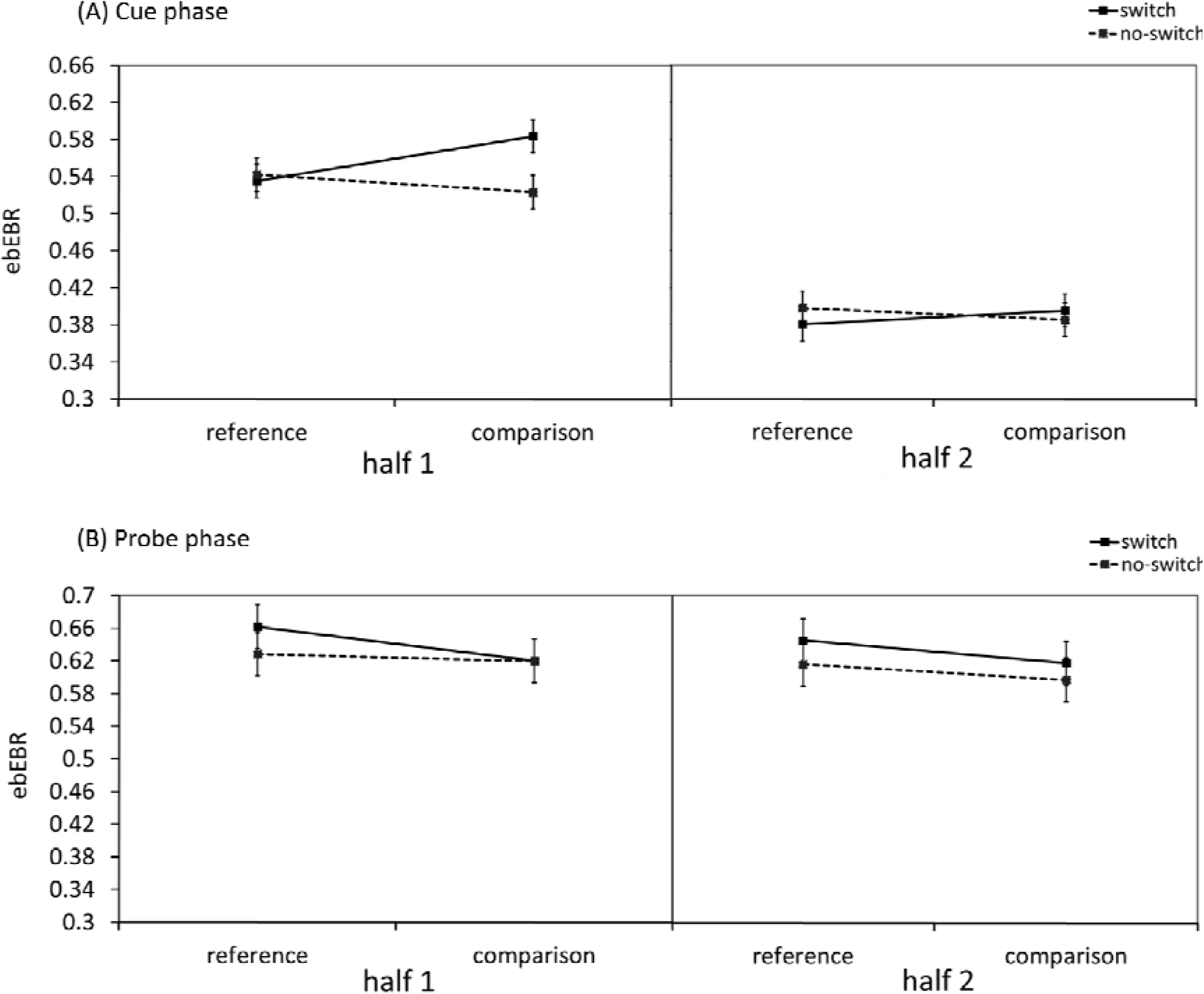
The ebEBR results of Experiment 3. The ebEBR as a function of Segment part, Trial Type and Switching is presented. The 4-sec segment length after (a) cue presentation and after (b) stimulus presentation was divided into two parts, each of 2 sec. In the cue-locked presentation, the switch cost was observed only in comparison trials and only in the first half of the segment. In the stimulus-locked presentation, the switch cost in the reference trials was observed in both segment parts. Error bars represent 95% confidence intervals.

## Probe phase analysis

The effect of stimulus presentation on ebEBR was examined in a three-way ANOVA with Segment Part (first half, second half), Trial-Type (reference, comparison) and Switching (switch, no-switch) as the within-subject independent variables (Figure 4b). The ebEBR pattern did not differ between the two segment parts. Neither the main effect of Segment Part nor any interaction that included this factor were significant, all Fs<1. The main effect of Trial-Type, *F(1,20)*=*11.50, MSe*=*.0021, p*=*.003, η_p_^2^*=*.36,* was significant, reflecting a higher ebEBR in reference trials than in comparison trials. Additionally, the ebEBR was significantly higher in switch trials than in no-switch trials, *F(1,20)*=*5.75, MSe*=*.0032, p*=*.03, η_p_^2^*=*.22,* as in the original analysis. The two-way interaction between Trial-Type and Switching was only marginally significant, *F(1,20)*=*3.91, MSe*=*.0012, p*=*.06, η_p_^2^*=*.16.*

Because the cue phase analysis revealed a significant switching effect only in comparison trials and not in reference trials, we decided to decompose the two-way interaction of Trial-Type and Switching to its simple effects. This analysis showed a mirror image of the two-way interaction in the cue phase analysis. A significant switching effect was only observed in reference trials, *F(1,20)*=*11.96, MSe*=*.001, p*=*.002, η_p_^2^*=*.37,* but not in comparison trials, *F(1,20)*=*.74, MSe*=*.001, p*=*.40, η_p_^2^*=*.04.*

## Discussion

In this study, we demonstrated that the ebEBR, an inexpensive, non-invasive and easy-to-use measure that presumably reflects striatal dopamine activity ^26,30,40,49^, can be used to track changes in demands on WM during task performance. The ebEBR results in all three experiments demonstrated that the ebEBR follows changes in WM demands with a resolution of a few seconds, thereby extending previous studies that reported a relationship between performance on cognitive tasks and the more familiar sEBR measure, which is recorded over several minutes during resting conditions^37–39^ (see additional analysis of the ebERB measure in the Supplementary Materials online). Specifically, the ebEBR recorded over 4 and even 2 seconds increased in conditions of the reference-back task, which presumably relies on fronto-striatal DA^9^, and mirrored the behavioral results (see Supplementary Fig. S4-S6 online). Specifically, reference trials, which required updating of WM, and trials that required switching the state of the gate led to an increase in the ebEBR, in line with our predictions that were inspired by the PBWM model^9,10^, with the exception that the reported results suggest that gate closing might also be DA-dependent. The reported findings may suggest that gate closing is not automatic but rather, similar to gate opening, might involve a phasic DA response and that perhaps the default state of the gate is not always closed but rather might be dependent on context^51^. More generally, these findings suggest that the ebEBR method^42,49^ is a viable method to track DA-based changes in WM demands during task performance.

Finally, in Experiment 3, the on-line measure of ebEBR enabled testing of whether contextual information provided by a cue can be used to prepare for gate switching in advance. The ebEBR analysis revealed a dissociation in the preparation time between switching to reference trials (updating mode) and switching to comparison trials (maintenance mode). Specifically, the context cue led to an increase in the ebEBR before the stimulus was presented only when switching to comparison trials. When switching to reference trials, this increase in ebEBR was, however, observed only after the stimulus was presented. These novel findings may suggest that gate closing is more proactive than gate opening. A possible explanation for this is that WM updating does not always take place following gate opening, but rather only in trials where the stimulus is different than the previous reference. Accordingly, it might be more beneficial to use a wait-and-see strategy in those trials, and only open the gate to WM after the stimulus identity is revealed. Thus, contextual information (in the form of a red frame) only provides the information that updating is possible, which may not be sufficient to trigger gate opening. By contrast, switching to a maintenance mode can be initiated by contextual information (in the form of a blue frame) because it provides the information that no updating will be required and, thus, may trigger gate closing without the stimulus information. The process of gate opening after stimulus presentation may nevertheless be facilitated by knowledge of an upcoming switch; as shown in Experiment 3, the gate opening-related increase in the ebEBR was already observed in the first 2-sec after stimulus onset, whereas in Experiment 2, the switching effect was detectable only in the last 2-sec of the segment. Evidence for preparation is also indicated by the significantly reduced switching effect in RT compared to that in Experiments 1 and 2 (see Supplementary Fig. S4-S6 online). Indeed, studies have shown that the cognitive system can at least partially be reconfigured in preparation for a switch in the task set^54,55^.

To conclude, our findings from the three experiments confirm that the ebEBR can be used as a measure of cognitive control over WM. The significant switching effect observed in the cue phase analysis illustrates that the ebEBR can dynamically track cognitive control processes that are not tied to any response-related processes and, thereby, provide information that cannot be extracted from RT patterns alone. More generally, the reported ebEBR results provide further support for the notion that the ebEBR can be used to track changes in cognitive functions, which are based on striatal DA activity, during task performance.

We acknowledge that although the EBR is an established physiological indicator of striatal DA activity^26–30^, EBR is still an indirect measure of striatal DA activity. Future studies that combine ebEBR measurements during task performance with pharmacological manipulations and pupil measurement and/or neuroimaging are necessary to establish the neurochemical and neural mechanisms underlying the observed ebEBR modulations. For example, determining to what extent the ebEB effects that we have presented here are related to WM processes and phasic DA activity vs. non-specific processes, such as arousal^45,56^, would be helpful. Nevertheless, our findings indicate that the ebEBR provides a viable online measure that can aid in investigating DA-based cognitive processes in populations in which pharmacological alteration of DA is not feasible or sensible, such as in infants, elderly and recreational cocaine users. As such, this method may enhance our understanding of the mechanisms underlying cognitive dysfunction in psychiatric disorders characterized by DA abnormalities, such as Parkinson’s disease, schizophrenia and ADHD.

## Method

### Participants

Twenty undergraduate students from Ben-Gurion University of the Negev participated in Experiment 1 (6 males; age: M=24.45 years, S.D.=2.06) and in Experiment 2 (3 males; age: M=24.15 years, S.D.=.91). Twenty-two undergraduate students from Ben-Gurion University of the Negev participated in Experiment 3 (4 males; age: M=22.81 years, S.D.=1.43). One participant was removed from the analysis of Experiment 1 due to low accuracy in the reference-back task (<50% in some of the conditions), and one participant was removed from the analysis of Experiment 3 due to noisy recording of the eye blinks. Participants were either paid for their participation or received partial fulfillment of course requirements. Informed consent was obtained from all participants in accordance with the Department at Ben-Gurion University of the Negev. The experimental protocol was approved by the Ethics Committee of the Department of Psychology at Ben-Gurion University of the Negev in accordance with American Psychological Association guidelines.

### Stimuli and Apparatus

Stimuli presentation and behavioral data collection were performed using E-Prime v2.0 (Psychology Software Tools, Pittsburgh, PA). The stimuli were the letters “X” and “O”, in font size 36, presented in black against a light gray background within a red or a blue frame. Responses were collected using a serial response box. Note that a stimulus set of only 2 stimuli was chosen for two reasons. First, it maximizes the control required to answer correctly. The smaller the stimulus set, the larger the probability that the present stimulus was presented in the previous trials, leading to a strong familiarity signal in each trial. Control is required to overcome familiarity and base the response on recollection, which addresses the precise context of the stimulus in WM. The most extreme case is with only 2 stimuli. Second, this design facilitates a balanced manipulation of stimuli and conditions as “same” and “different” responses are equally probable and, thus, so are the conditions preceding the response (match/no-update, mismatch/update).

### Procedure

Each trial started with a presentation of the stimulus “X” or “O” at the center of the screen (see below the exception in Experiment 3). The stimulus was presented in black inside either a red or a blue frame. After the response, a fixation screen was presented with three dots at the center of the display to maintain foveal perception of the participants. The reference-back task was composed of two trial-types: reference and comparison (see Fig. 1). The stimulus in each trial (an X or O) was selected at random. The first trial in a block was always presented in the reference color and did not require a response. In each of the following trials, the participants had to indicate whether the stimulus was the same as or different than the *most recent reference trial.* “Same” and “different” responses were indicated using the right and left index fingers, respectively, using a serial response box. Participants were instructed to be as fast as possible.

In Experiments 1 and 2 we used a fixed alternating-runs order of trial-types composed of 4 trials of each condition in a row (see Fig. 1). In Experiment 3, the sequence length of each trial-type (reference, comparison) was random. The probability of a switch between trial-types was 50% in each trial. A cue indicating the trial-type was added before the stimulus presentation. Each trial was initiated with a cue, namely, a red or blue empty frame, presented for 4-sec (see Fig. 5). In Experiment 1, the stimulus was presented until a response was provided. After the response, a fixation screen was presented for a 2-sec inter-trial interval (ITI). In Experiments 2 and 3, the stimulus was presented inside the frame until a response was given or until 3-sec had elapsed. After the response, a fixation screen was presented until 4.5-sec had elapsed from the stimulus presentation (i.e., the ITI was 4.5-sec minus the RT in each trial). Experiments 1 and 2 comprised 12 blocks, including 48 trials each, preceded by 2 practice blocks. Each block was followed by a break phase that was not limited in time but was rather controlled by the participants. In half of the blocks, reference trials were indicated by a red frame and comparison trials by a blue frame and vice versa in the remaining blocks (with a counterbalanced order). Experiment 3 comprised 8 blocks, including 40 trials each. Participants completed one practice block before they began the experiment. The colors used to indicate the trial-types were counterbalanced *between* participants.

**Figure 5.**
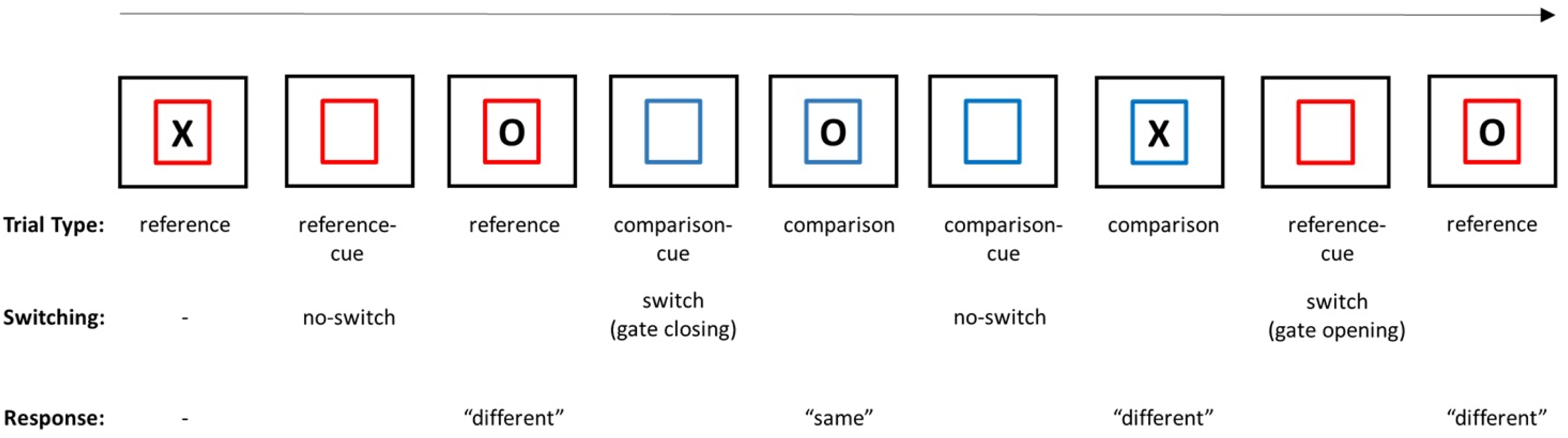
The cued reference-back task. The sequence length of each trial type was random. The fixation display was presented after the response until 4.5 sec had elapsed from the stimulus presentation. Thus, the inter-trial interval (ITI) varied as a function of the response time. The cue was presented before the stimulus as an empty colored frame for 4 sec. The state of the gate and the correct response for each trial are indicated below each stimulus display.

Baseline sEBR was also measured at the beginning of the experiment before the reference-back task was introduced. In Experiment 1, the sEBR was measured for 4 minutes while participants viewed a silent short video of a waterfall. In Experiments 2 and 3, sEBR was measured for 5 minutes while participants viewed a fixation display. The only instructions given for this recording were to view the display silently. The by-condition correlations between sEBR and the other dependent variables are presented in the Supplementary Materials online.

### Eye blink recording and analysis

Eye blinks were recorded using a BioSemi Active Two system. Two external electrodes were placed above and below the right eye. Because the sEBR increases in the evening^57^, participants were tested between 10 am and 5 pm. In addition, participants were asked to avoid alcohol, nicotine, and caffeine consumption prior to the experiment and to sleep well the night before the recording. During recordings, participants did not wear contact lenses. Importantly, they were not instructed in any manner about blinking. After recording, participants were asked what they thought we measured, and none of them suspected that eye blinks were recorded.

The data were acquired using a 0.01–100-Hz bandpass filter and offline filtered using a 1 Hz high-pass and 40 Hz low-pass filter (IIR Butterworth filters, attenuation slope of 12 dB/octave). The sampling rate was 256 Hz. The signal was digitized using a 24-bit A/D converter. The electrooculography (EOG) was segmented between stimulus onset and 2-sec post-probe in Experiment 1 and 4-sec post-probe in Experiments 2 and 3, using EEGLAB^58^. In Experiment 3, the EOG was also segmented between cue onset and 4-sec post-cue. Eye blink detection was performed using a MATLAB code based on the VEOG channel, created from the difference between the electrode above and under the eye, followed by manual inspection. Then, the ebEBR per second was calculated for each condition.

## Acknowledgments

This research was funded by the Israel Science Foundation (grant #458/14) awarded to Y.K.

## Author contributions

All authors participated in the design of the study, the interpretation and writing of the final manuscript. R.R-L conducted the experiments and analyzed the data.

